# Rab9 depletion enhances human adenovirus type 26 transduction efficiency through increased internalization and reduced late endosomal/lysosomal retention

**DOI:** 10.64898/2026.02.24.707667

**Authors:** Isabela Drašković, Davor Nestić, Lucija Lulić-Horvat, Jelena Martinčić, Mario Stojanović, Gabriela N. Condezo, Klara Kašnar, Carmen San Martín, Jerome Custers, Dragomira Majhen

**Affiliations:** Laboratory for Cell Biology and Signalling, Division of Molecular Biology, Ruđer Bošković Institute, Zagreb, Croatia; Department of Macromolecular Structures, Centro Nacional de Biotecnología (CNB-CSIC), Madrid, Spain; Janssen Vaccines and Prevention B.V., Leiden, The Netherlands

## Abstract

Understanding intracellular trafficking is critical for viral pathogenesis and for the rational design of viral vectors, as endocytic routing determines genome release, immune sensing, and overall transduction efficiency. Human adenovirus type 26 (HAdV-D26) presents a promising platform for vector design due to its low preexisting immunity, potent immune stimulation, scalable production, and versatile genetic engineering capacity. Although increasingly significant, the fundamental mechanisms governing HAdV-D26 intracellular trafficking are still not fully understood. Our study demonstrates that compared to well-described human adenovirus type 5 (HAdV-C5), HAdV-D26 undergoes prolonged intracellular trafficking, transiently localizing to early endosomes before residing in late endosomes/lysosomes for up to four hours post-infection. Inhibition of lysosomal acidification modestly enhances HAdV-D26 transduction efficiency, whereas blocking transport from early to late endosomes/lysosomes does not. Strikingly, Rab9 knockdown reduces HAdV-D26 late endosomal/lysosomal localization while increasing both virus internalization and genome delivery of HAdV-D26. These findings indicate that late endosomal sorting pathways actively influence HAdV-D26 infection outcomes. Together, our results provide new mechanistic insight into HAdV-D26 intracellular trafficking, highlight serotype-specific differences in adenovirus entry pathways, and identify endosomal trafficking steps that may be targeted to improve adenoviral vector performance.

**Author summary:** Human adenovirus type 26 is an important platform for vaccines and gene therapy due to its low preexisting immunity and versatility as a vector. Despite its growing importance, how HAdV-D26 enters and moves within host cells is still not well understood. In this study, we show that compared to HAdV-C5, HAdV-D26 exhibits prolonged intracellular trafficking, transiently passing through early endosomes and accumulating in late endosomes and lysosomes for several hours after entry. We further identify the host trafficking protein Rab9 as an important regulator of HAdV-D26 infection, as altering Rab9 levels changes viral internalization, late endosomal localization, and delivery of the viral genome. These findings reveal previously unrecognized mechanisms that control adenovirus intracellular trafficking and demonstrate that different adenovirus types use distinct cellular entry routes. Understanding these pathways provides insights that may guide the development of more effective and better-controlled adenovirus-based vaccines and gene delivery vectors.

## Introduction

The benefits of human adenovirus (HAdV) vectors have been widely recognized in gene therapy, vaccination, and cancer treatment due to their high transduction efficiency, ability to infect both dividing and non-dividing cells, and capacity for large-scale production (1). Traditionally, adenovirus type 5 (HAdV-C5) has been the vector of choice; however, recent research has increasingly focused on alternative adenovirus types. This shift is primarily driven by the prevalence of pre-existing immunity to HAdV-C5 in the general population (2), which can significantly diminish its therapeutic efficacy.

Human adenovirus type 26 (HAdV-D26) has emerged as a particularly promising vector for vaccination, largely due to its low seroprevalence and strong immunogenicity. The versatility of HAdV-D26 in addressing global health emergencies was demonstrated by regulatory approval as a vaccine vector for both COVID-19 (Ad26.COV2.S) and Ebola virus disease (Ad26.ZEBOV). These approvals highlight HAdV-D26 validated efficacy in inducing robust immune responses through heterologous prime-boost regimens (3). Despite its growing importance, the fundamental mechanisms underlying HAdV-D26 intracellular trafficking remain incompletely understood. General knowledge on adenovirus biology indicates that following receptor binding, the virus enters the cell through endocytosis, and upon liberation from endosomes, it travels along microtubules toward the nucleus for nuclear import and initiation of the viral replication cycle (4). Recent studies indicate that the successful cell entry and transduction efficiency of HAdV-D26 in epithelial cells is strongly dependent on the expression of αvβ3 integrin, which facilitates virus internalization (5). While αvβ3 integrin-mediated HAdV-D26 infection involves dynamin-2 and is caveolin-1-dependent, HAdV-D26 infection of epithelial cells with low expression of αvβ3 integrin involves dynamin-2 and clathrin, and is caveolin-1-independent. Downregulation of clathrin resulted in increased HAdV-D26 infection due to elevated expression of αvβ3 integrin, whilst inhibition of clathrin-coated pits disabled HAdV-D26 transport through the cytoplasm, indicating that in cells with low expression of αvβ3 integrin, HAdV-D26 infection could be clathrin-mediated (6). Data regarding HAdV-D26 intracellular trafficking are limited, with only one study reporting that HAdV-D26 accumulates in the late endosomal compartment more extensively than HAdV-C5 at 2 to 8 h following infection (7).

After internalization, adenoviruses reside within primary endocytic vesicles, endosomes, from where they need to escape to reach the nucleus. Early endosome escape in case of HAdV-C5 is mediated by the viral membrane lytic protein VI (8), however, a discussion regarding the role of acidic pH in endosomal membrane penetration of HAdV-C2 and -C5 is still active (9). Different adenovirus types exhibit different trafficking pathways within the host cell and reside within different endosomes. Species B types accumulate in lysosomes, whereas species C types traffic rapidly to the nuclear envelope, revealing different kinetics of endosomal escape. The optimum pH for membrane lysis matches that of early sorting endosomes in adenoviruses of species C, and that of late endosomes or lysosomes for species B (10).

Endosomes continuously remodel and mature by lowering the pH inside their lumen, moving toward the perinuclear region, and altering their shape. Several proteins are dynamically exchanged on the endosomal membrane, notably members of the Rab family of small GTPases. The human genome encodes over 60 distinct Rab proteins, each playing specialized roles in membrane trafficking. Among those, Rab5 is crucial for the biogenesis and function of early endosomes, orchestrating their formation and maturation, while Rab7 and Rab9 are primarily involved in regulating the activities of late endosomes, ensuring proper cargo sorting and transport within the endocytic pathway (11). In the context of adenovirus cell entry, the role of Rab5 and Rab7 has been reported in the regulation of adenovirus endocytosis. Specifically, Rab5 controls the endocytosis and trafficking of the escape-defective, temperature-sensitive HAdV-C2_ts1 (Ad2-ts1) mutant to late endosomes, but does not influence the infection process of wild-type HAdV-C2. On the other hand, sorting of HAdV-C2_ts1 to late endosomes was independent of Rab7 while HAdV-C2 and - C5 infection was independent of EEA1, a marker of early endosomes (12). While HAdV-C2_ts1 is degraded in lysosomes (13), HAdV-B7 retention in a late endosomal compartment did not cause a loss in infection efficiency. Tracking trafficking of HAdV-B7 to late endosomes revealed partial co-localization with late endosomal and lysosomal marker proteins, including Rab7, mannose-6-phosphate receptor, and LAMP1 (14). So far, no studies have reported Rab9 implication in the adenovirus infection pathway.

Understanding adenovirus intracellular trafficking is essential because efficient therapeutic gene delivery depends on successful cellular entry, endosomal escape, and nuclear import. Insights into these processes help improve vector efficacy, safety, and specificity, enhancing clinical outcomes. Notably, HAdV-D26 uses αvβ3 integrin for cell entry, directing it toward caveolin-mediated endocytosis (5, 6), however, HAdV-D26 intracellular transport mechanisms, particularly the functional role of endosomes as well as the role of endolysosomal regulators like Rab GTPases, remain poorly characterized.

In this work, we employed quantitative analyses to determine which specific endosomal compartments HAdV-D26 associates with during its intracellular trafficking, as well as the role of endosomes in HAdV-D26 transduction efficiency. Our study reveals that, compared to HAdV-C5, HAdV-D26 exhibits prolonged intracellular trafficking resulting in reduced genome delivery to the nucleus, including transient early endosome localization and extended lysosomal residence. We report that Rab proteins, critical coordinators of vesicular transport and organelle maturation, directly influence HAdV-D26 trafficking dynamics. Namely, Rab9 knockdown increases HAdV-D26 transduction efficiency by increasing both virus internalization and genome delivery to the host nucleus, and decreasing HAdV-D26 retention in late endosomes/lysosomes, suggesting a previously unrecognized dependency on the Rab9-mediated pathway for efficient transduction of HAdV-D26. These findings provide mechanistic insights into post-entry events that could inform strategies to optimize HAdV-D26 vector efficiency aimed at therapeutic applications.

## Results

### HAdV-D26 displays prolonged intracellular trafficking, resides in different cellular compartments during the first hour of internalization, and exhibits aberrant nuclear entry compared to HAdV-C5 in epithelial cells

Currently, only one study is addressing the intracellular trafficking of HAdV-D26, indicating that this virus accumulates in the late endosomal compartment (7). To investigate this behavior in more detail, we performed confocal microscopy and transmission electron microscopy (TEM) analysis of HAdV-D26 intracellular trafficking in A549 cells. In contrast to well-characterized intracellular trafficking of HAdV-C5, which typically reaches the nucleus within approximately 60 min *in vitro* (Fig. S1), most HAdV-D26 particles remained dispersed throughout the cytoplasm 60 min after transduction (Fig. 1A). Even 240 min p.i. (post-infection), there is no significant accumulation of HAdV-D26 near the nucleus. However, most viral particles appear grouped on one side in the wider perinuclear area (Fig. 1A). To determine whether HAdV-D26 is free in the cytosol or associated with membrane-bound organelles, we investigated its intracellular trafficking in A549 cells 60 min after internalization using TEM. HAdV-D26 was observed at the plasma membrane, in several distinct cellular compartments including endosomes and degradative compartments, as well as free in the cytosol (Fig. 1B). Quantitative analysis of viral localization revealed that the majority of the HAdV-D26 particles were found at the cell membrane, within degradative compartment and in the cytosol, with nearly 40% localized in the degradative compartment (Fig. 1C). These results suggest that after internalization, a distinct intracellular trafficking pattern of HAdV-D26 in A549 cells may contribute to its lower transduction efficiency compared to HAdV-C5 (5).

**Figure 1.**
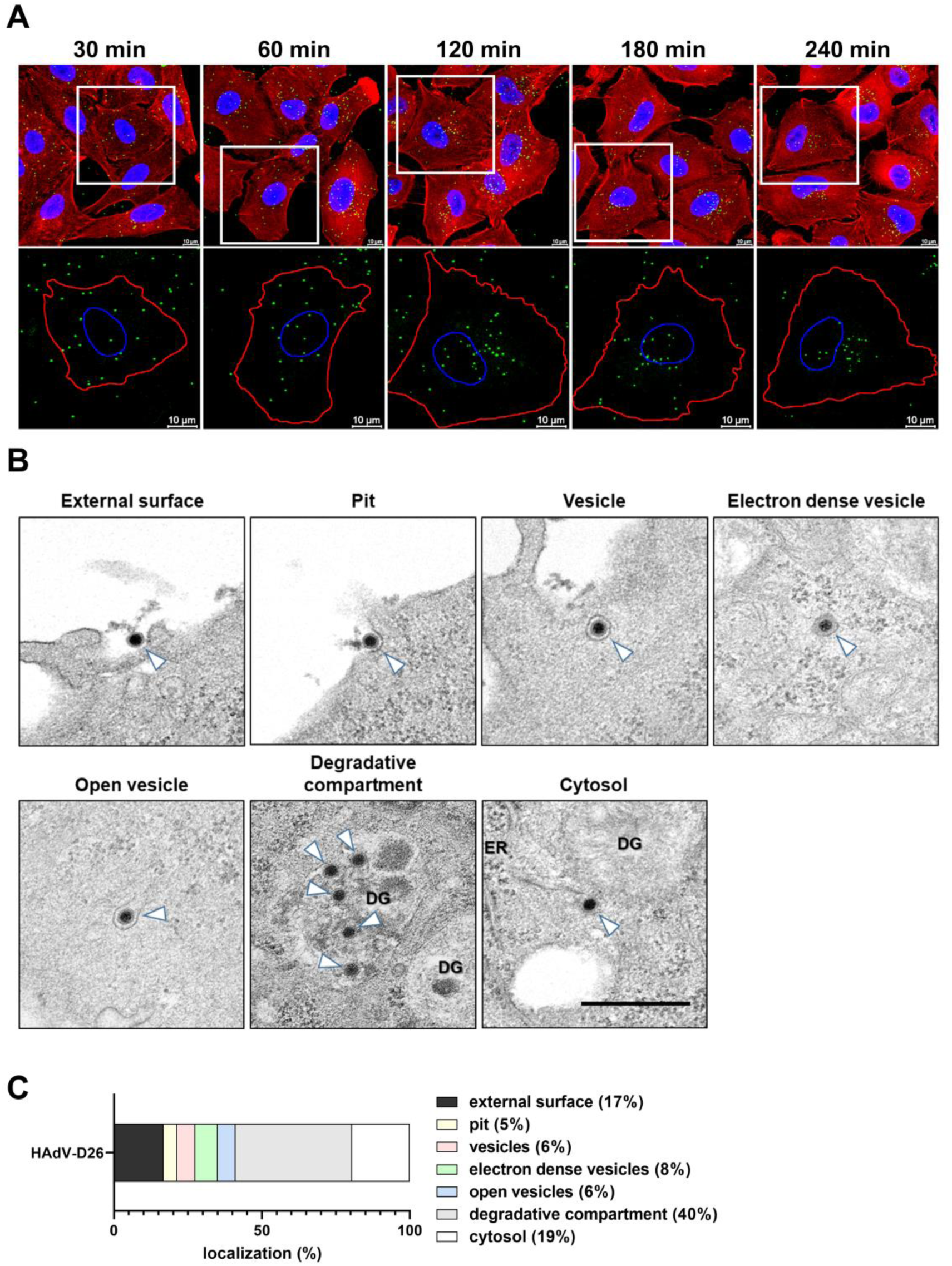
Intracellular trafficking of HAdV-D26 in epithelial cells. (A) Intracellular trafficking of HAdV-D26 in A549 cells observed by confocal microscopy. Cells were incubated on ice with fluorescently labeled HAdV-D26 (green; MOI 10^5^ vp/cell) for 45 min, then 10 min at 37°C, followed by washing with fresh medium to remove unbound viruses. After additional incubation at 37°C for the indicated times, cells were fixed and stained with phalloidin-AF555 (red) and DAPI (blue). Representative confocal images are shown (scale bar = 10 μm). The lower panels show areas selected from the upper panels at larger magnification, with marked edges of the nucleus (blue, based on DAPI) and cell (red, based on phalloidin). (B) HAdV-D26 resides in different cellular compartments during the first hour of internalization and intracellular trafficking. Representative TEM images of intracellular trafficking of HAdV-D26 in A549 cells. Cells were incubated on ice with HAdV-D26 (MOI 2×10^5^ vp/cell) for 45 min, then 10 min at 37°C, followed by washing with fresh medium to remove unbound viruses, and then incubated for 1 h at 37°C. Cells were then fixed and prepared for TEM. Scale bar = 500 nm. Endoplasmic reticulum (ER); Degradative Compartment (DG). White arrowheads denote virus particle. (C) Quantification of HAdV-D26 localization in different cellular compartments in A549 cells shown in (B). Number of cells: 56; number of viral particles: 262 total viral particles.

To gain better insight into the kinetics of genome delivery by HAdV-D26, we quantified its nuclear entry in A549 cells using qPCR with primers specific for the CMV promoter present in the viral genome. HAdV-D26 DNA was quantified at 0 h, 1 h, 4 h, and 8 h p.i. from both total cell lysate and isolated nuclei. HAdV-D26 genome delivery to nuclei was calculated as the ratio of HAdV-D26 DNA in a nuclear fraction to HAdV-D26 DNA in total cell lysate. Most of the HAdV-D26 DNA was delivered to the nucleus within 1 h (Fig. 2A); however, surprisingly, in the subsequent time points, we observed decreased HAdV-D26 DNA delivery to the nucleus, which would indicate DNA degradation or some issues with HAdV-D26 DNA nuclear import. These data are in accordance with the modest transduction efficiency of HAdV-D26 compared to HAdV-C5 and - B35 (5). Contrary to HAdV-D26, HAdV-C5 genome delivery to the nucleus slightly increased over time (Fig. 2A).

**Figure 2.**
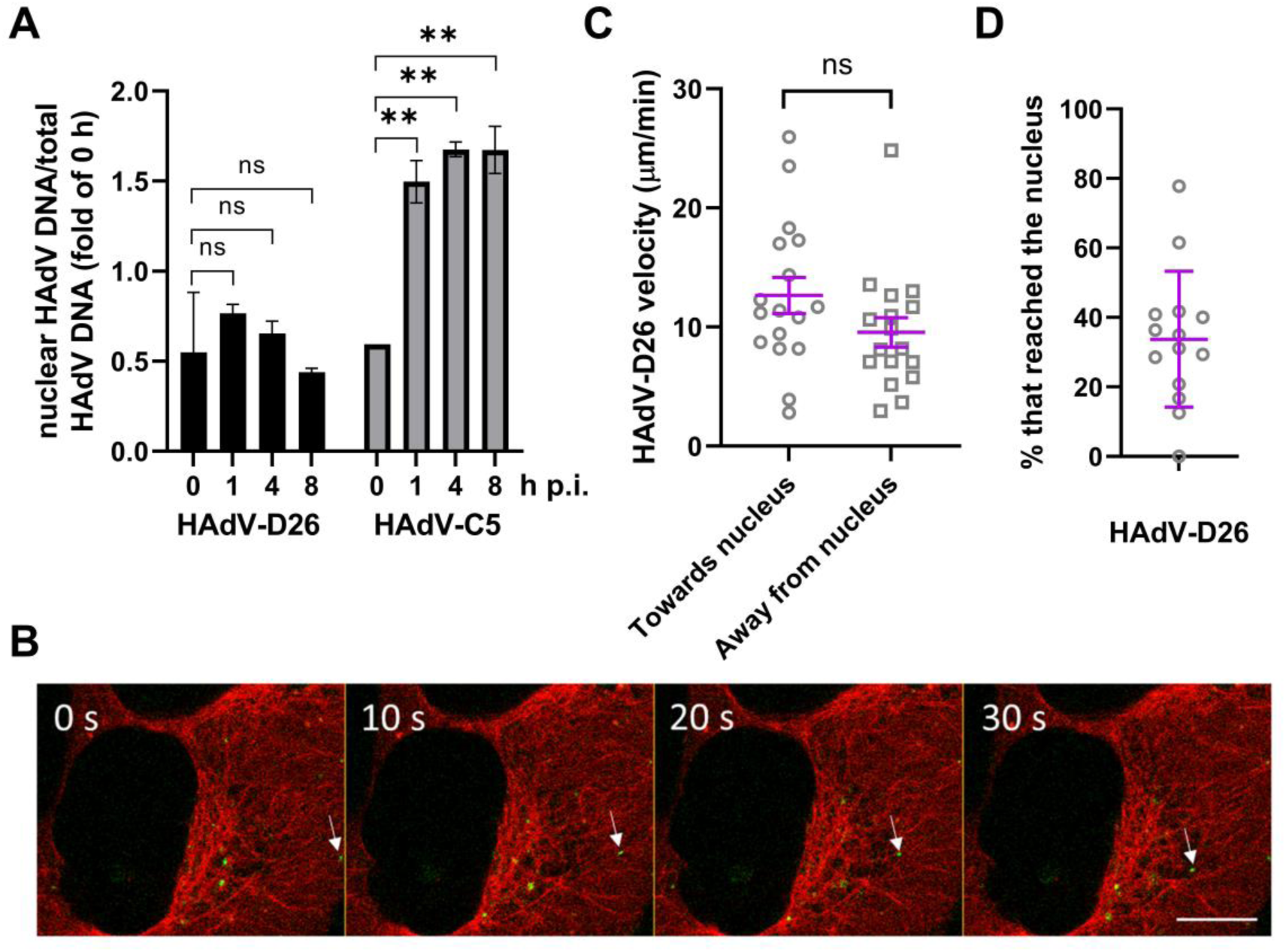
HAdV-D26 shows impaired nuclear entry dynamics. A) HAdV-D26 genome delivery into the host nucleus. A549 cells were infected with HAdV-C5 or HAdV-D26, MOI 10^4^ vp/cell for 45 min on ice to ensure uniform virus entry into the cell. Five min after incubation at 37°C unbound viruses were removed and fresh medium was added, followed by incubation at 37°C. Cells were harvested at defined time intervals. DNA was extracted from the total cells or isolated nuclei. Viral and cellular DNA were quantified by qPCR using primers for the CMV promoter present in the viral DNA or the cellular GAPDH gene. Results are presented as a ratio of viral DNA measured in the nuclear fraction to the total fraction, normalized to the value for 0 h. Data are presented as mean values ± SD from three experiments. B) HAdV-D26 spatial distribution in U2OS mCherry-tubulin cells. U2OS cell with stable expression of mCherry-tubulin (red) infected with fluorescently labeled HAdV-D26 (green), MOI 5×10^4^ vp/cell, representing fast movement of HAdV-D26 (white arrow) towards the nucleus. Scale bar represents 10 µm. C) Average velocity of rapid HAdV-D26 movements directed towards or away from the nucleus in U2OS cells (p = 0.12, unpaired t- test, 13 cells, 17 viral particles in both directions). D) Percentage of HAdV-D26 viral particles that reached nuclear surface of U2OS cells within an hour after infection (total of 206 viral particles from 14 cells). **, P < 0.01

After endosomal escape, adenoviruses use motor proteins for bidirectional trafficking on microtubules. In non-polarized cells, dynein-based minus-end directed transport prevails over the plus-end directed kinesin-based transport, leading to the virus enrichment near the centrosome proximal to the nucleus (15). Therefore, we tracked the movements of internalized fluorescently labeled HAdV-D26 and measured the velocity of its directed motions towards or away from the nuclear surface. HAdV-D26 virions usually followed the path of microtubule fibers (Fig 2B) and average HAdV-D26 velocity was 12 ± 6 µm/min towards the nucleus and 9 ± 5 µm/min away from the nucleus (Fig. 2C). This is within the values reported for HAdV-C5 (1.2 to 24 µm/min) (16), however on the lower end, indicating HAdV-D26 moves slower along microtubules than HAdV-C5. Although HAdV-D26 exhibits movements towards the nucleus, only an average of 34 % viral particles among those with rapid movements reached the nuclear surface within one hour from infection (Fig. 2D).

### HAdV-D26 transiently visits the early endosome and resides in lysosomes up to 240 min post-infection

It is known from the literature that HAdV-C2 and -C5 visit early endosomes but are not transported to late endosomes (12). In order to identify cytosolic HAdV-D26 particles and distinguish them from those inside membranous compartments, we carried out streptolysin O (SLO) assays. SLO-mediated perforation of the plasma membrane is used to introduce antibodies into the cytosol, and accessibility of Alexa-488-labeled virus to anti-Alexa-488 antibodies distinguishes cytosolic from endosomal viruses in the permeabilized cells (17). As shown in figure 3A, at 45 min p.i. only 44% of HAdV-D26 particles are positive for anti-Alexa-488 antibody representing cytosolic particles, an intermediate value between those of HAdV-C5 and the entry-defective HAdV-C2_ts1. These data allowed us to conclude that the majority of HAdV-D26 particles remain longer within some endosomal compartment, however we did not know which one. Thus we decided to revisit HAdV-D26 localization within early and late endosomes. For that purpose, we observed co-localization of HAdV-D26 and EEA1, a marker of early endosomes, during 120 min p.i., as well as co-localization of HAdV-D26 and LAMP1, a marker of late endosomes and lysosomes, up to 240 min p.i. (Fig. 3B). Our confocal microscopy data indicated that 30 min p.i. about 63% of the incoming HAdV-D26 co-localized with EEA1 (Fig. 3C). At later time points, co-localization with EEA1 decreased. However, even at 120 min p.i. about 27% of HAdV-D26 was still localized in early endosomes. At 30 min p.i. about 30% of HAdV-D26 co-localizes with LAPM1, indicating that already at early points of infection HAdV-D26 is found in late endosomes or lysosomes. At 120 min p.i., nearly 80% of HAdV-D26 remains within early or late endosomes/lysosomes, and even at 240 min p.i., over 50% still co-localizes with the late endosomal/lysosomal marker LAMP1 (Fig. 3C). Retention of HAdV-D26 in late endosomes/lysosomes is in line with our TEM results where we observed HAdV-D26 in a degradative compartment.

**Figure 3.**
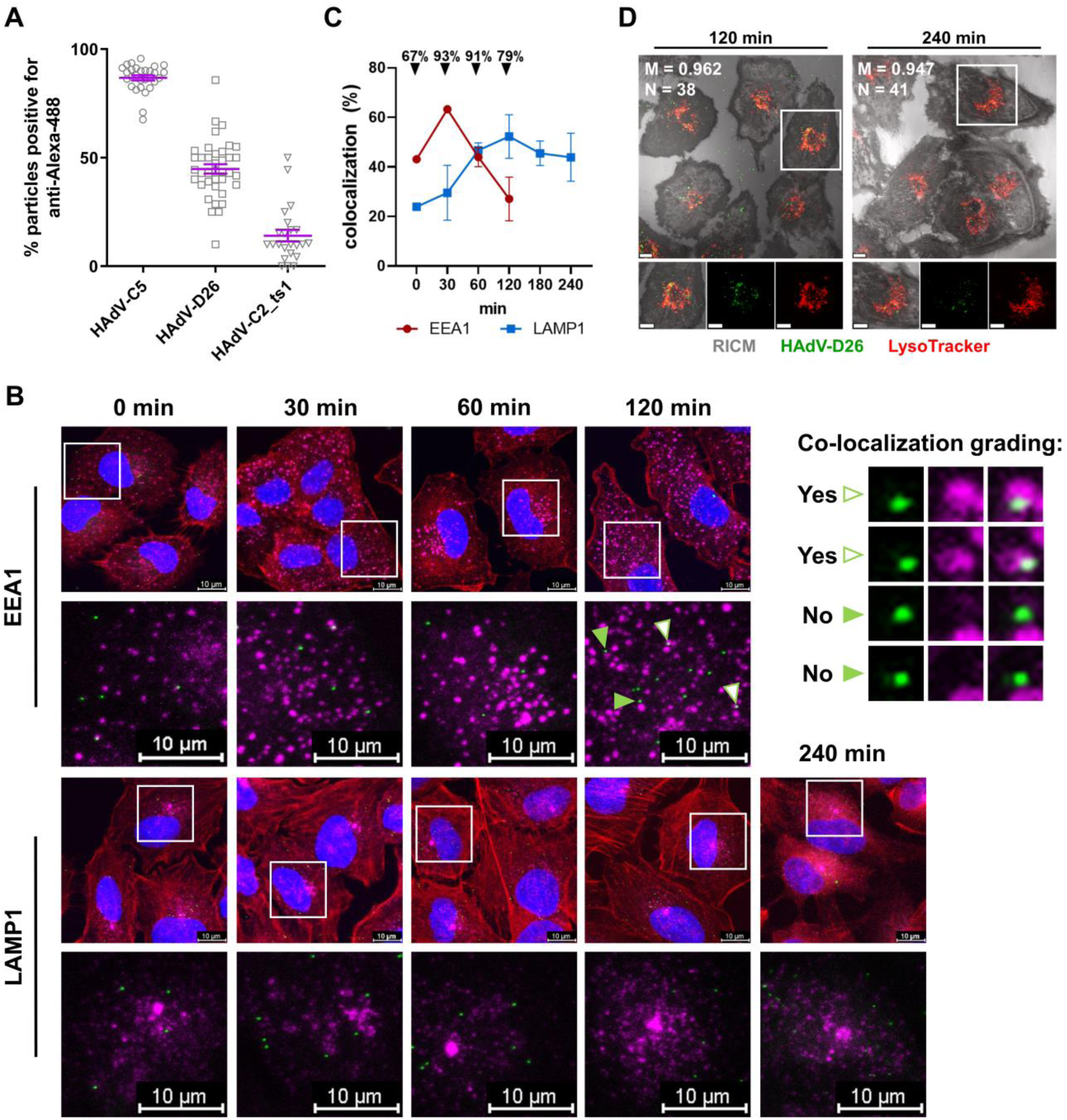
HAdV-D26 shows localization within lysosomes as long as 240 min p.i. A) Membrane penetration of HAdV-C5, HAdV-D26 and HAdV-C2_ts1 in A549 cells measured by SLO penetration assay 45 min p.i. Quantification of cytosolic virus particles in cells infected with Alexa-488-labeled HAdVs. Intact cells were treated with SLO and then incubated with anti-Alexa-488 antibody which was detected by secondary Alexa-633-conjugated antibody. The plot shows the percentages of virus particles positive for anti-Alexa-488 antibody. One dot represents one cell. Data are presented as mean values ± SEM. B) Localization of HAdV-D26 within early and late endosomes/lysosomes upon internalization. Representative confocal images are shown. Cells were incubated on ice with fluorescently labeled HAdV-D26 (green; MOI 10^5^ vp/cell) for 45 min, then 10 min at 37°C, followed by washing with fresh medium to remove unbound viruses. After additional incubation at 37°C for the indicated times, cells were fixed and stained with an antibody against EEA1 or LAMP1 (magenta), phalloidin-AF555 (red), and DAPI (blue). The upper right panel shows examples of co-localization grading: white arrows indicate co-localization, whereas green arrows indicate lack of co-localization. C) Quantification of co-localization of HAdV-D26 with either EEA1 or LAMP1 at different time points shown in (B). Percentages in black above the plot show the total amount of viruses found in any endosome (either EEA1 or LAMP1 positive). Data are presented as mean values ± SD from two independent experiments. D) Co-localization of HAdV-D26 with LysoTracker in live A549 cells. Cells were incubated on ice with fluorescently labeled HAdV-D26 (green) for 45 min, then 10 min at 37°C, followed by washing with fresh medium to remove unbound viruses, and then incubated at 37°C for the indicated times. In the last 30 min of incubation, Lysotracker Deep Red solution was added to the medium. After incubation, cells were washed with fresh medium and immediately observed by a confocal microscope. Reflection Interference Contrast Microscopy (RICM) is shown in grey. Representative confocal images are shown (scale bar = 10 μm). N denotes number of analyzed cells, M denoted Manders’ coefficient.

Since our TEM results showed significant accumulation of HAdV-D26 particles in structures consistent with degradative compartments, and we saw co-localization of HAdV-D26 with LAMP1-positive endosomes, we wanted to further identify the compartment where HAdV-D26 is retained. Thus, we performed co-localization of HAdV-D26 with LysoTracker, a marker for lysosomes, by live confocal microscopy. Significant co-localization between HAdV-D26 and LysoTracker was observed at 120 and 240 min p.i. (Fig. 3D), Manders’ coefficients are 0,962 and 0,947 respectively. These results further corroborate our data, indicating that HAdV-D26 resides within lysosomes up to 240 min p.i., which might influence HAdV-D26 transduction efficiency.

### Inhibiting lysosomal acidification, but not transport from early to late endosomes/lysosomes, modestly increases HAdV-D26 transduction efficiency

HAdV-D26 initially accumulates in early endosomes within 30 min p.i., then gradually shifts to late endosomes/lysosomes, where around 50% remains even at 240 min p.i. Since cargo directed to the lysosomes is typically destined for degradation, one can assume that prolonged residence of HAdV-D26 in late endosomes/lysosomes could lead to its degradation and may partly explain its low infectivity. To investigate whether disrupting lysosome function affects HAdV-D26 infectivity, we measured its transduction efficiency in cells treated with lysosomotropic agents such as bafilomycin A1 (BafA1), chloroquine, and ammonium chloride (NH_4_Cl). These inhibitors elevate intralysosomal pH, thereby impairing lysosomal function and disrupting autophagic protein degradation (18). Treatment with all three inhibitors modestly increased transduction efficiency of HAdV-D26, where only treatment with bafilomicin A1 was statistically significant (Fig. 4A). Nevertheless, the obtained data imply that at least some HAdV-D26 particles that trafficked to the late endosomes/lysosomes cannot complete the infection cycle.

**Figure 4.**
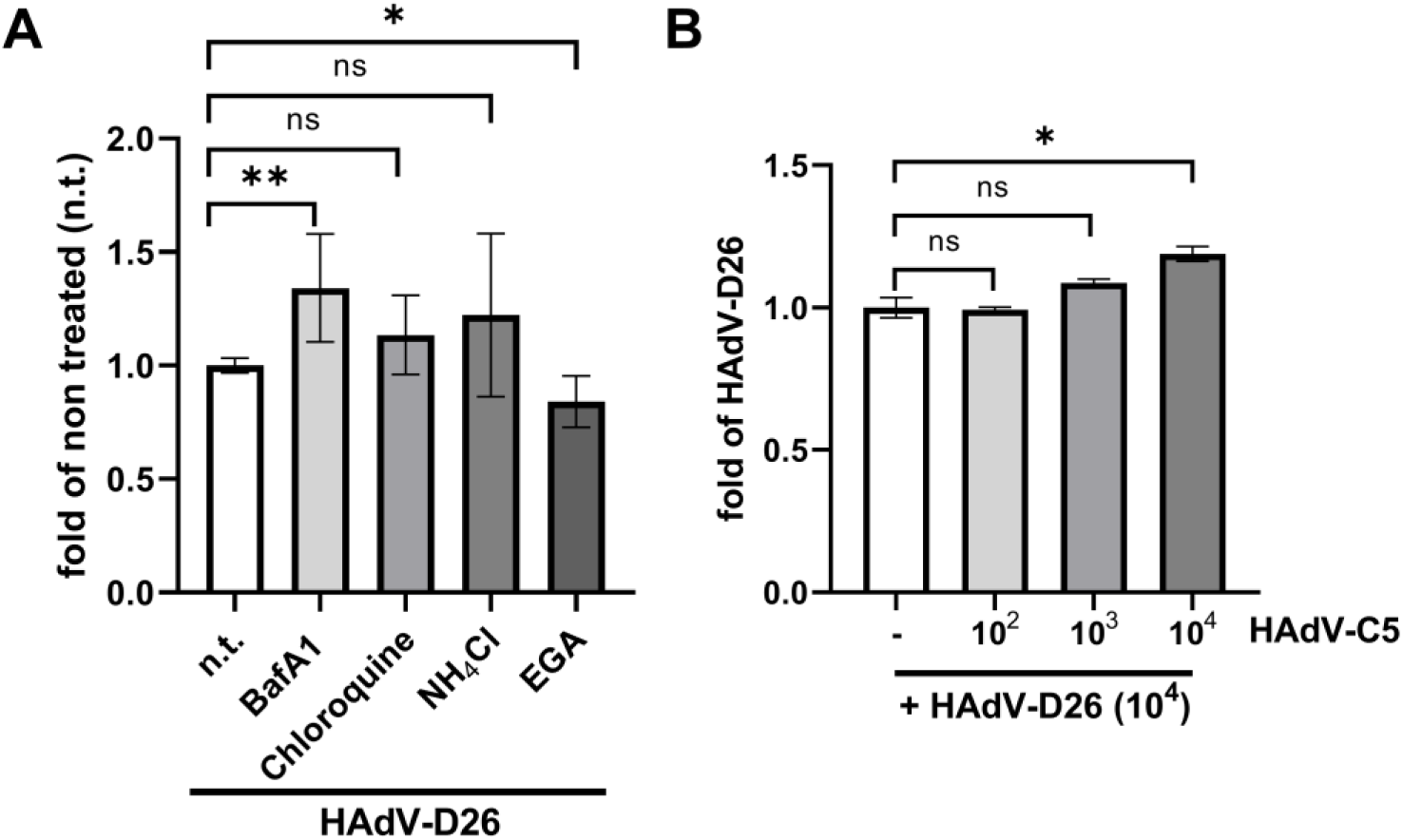
HAdV-D26 transduction efficiency is unaffected by endosomal pH modulation, inhibition of endosomal maturation, or co-infection with HAdV-C5. A) Increasing intralysosomal pH or inhibiting early-to-late endosome transition did not markedly affect HAdV-D26 transduction efficiency. Cells were pretreated with inhibitors (30 min, 37°C) - bafilomycin A1 (10 nM), chloroquine (50 µM), NH_4_Cl (5 mM) or EGA (15 µM) and subsequently incubated with HAdV-D26 for one hour (MOI 10^4^ vp/cell) in medium with the specific inhibitor, after which the medium was changed and fresh medium was added. After 24 h of incubation, the fluorescence intensity of eGFP encoded by HAdV-D26 and the fluorescence intensity of cellular DNA labeled by Hoechst was measured. The results were calculated as the ratio of eGFP/Hoechst values and normalized to infected cells that were not treated (n.t.). B) Co-infection with HAdV-C5 does not influence HAdV-D26 transduction efficiency. Cells were infected either with HAdV-D26 (MOI 10^4^ vp/cell) alone or with a mixture of HAdV-D26 (MOI 10^4^ vp/cell) and HAdV-C5 at different MOIs (10^2^, 10^3^, 10^4^ vp/cell) (1 h, 37°C), and then the medium was changed, and fresh medium was added. After 24 h of incubation, the fluorescence intensity of eGFP encoded by HAdV-D26 and the fluorescence intensity of cellular DNA labeled by Hoechst was measured. The results are presented as the ratio of eGFP/Hoechst values and normalized to cells infected with HAdV-D26. Data are presented as mean ± SD from two or three independent experiments in duplicates. *, P < 0.05; **, P < 0.01.

We also investigated whether HAdV-D26 transport from early endosomes to late endosomes/ lysosomes would affect its transduction efficiency. For that purpose, we used EGA, an inhibitor of endosomal trafficking that specifically blocks transport to LAMP1-positive compartments, thereby impacting lysosomal function without disrupting other recycling pathways (19). The late endosome trafficking inhibitor EGA only slightly decreased transduction efficiency with HAdV-D26 (Fig.4A), further corroborating our assumption that for efficient transduction, HAdV-D26 escapes from endosomes before reaching the lysosomes. Overall, these results indicate that trafficking from the early endosome to the lysosome is not essential for efficient HAdV-D26 transduction.

So far, our data suggest that HAdV-D26 might behave similarly to HAdV-C2_ts1, which does not efficiently escape from endosomes (20). On the other hand, HAdV-C2 is liberated from the early endosome rather quickly and very efficiently. When HAdV-C2_ts1 was co-infected with HAdV-C2, its normally increased co-localization in early endosomes was significantly reduced (12), implying that HAdV-C2 facilitates the release of HAdV-C2_ts1 from early endosomes. Based on a similar concept, we performed a co-infection experiment using HAdV-D26 (encoding an eGFP reporter gene) and HAdV-C5 (encoding a β-galactosidase reporter gene) at increasing multiplicities of infection (MOI), and assessed HAdV-D26 transduction efficiency by quantifying eGFP expression. Co-infection of HAdV-D26 with HAdV-C5 at a high MOI of 10⁴ increased HAdV-D26 transduction efficiency by only 18%, while co-infection at lower MOIs had no significant effect on HAdV-D26 transduction efficiency (Fig.4B). These data indicate that HAdV-C5 endosome penetration does not liberate HAdV-D26, i.e., most HAdV-D26 particles do not traffic through the same endosomal compartments as HAdV-C5.

### Functional Rab5, Rab7, and Rab9 are involved in HAdV-D26 transduction

After endocytosis, cargos are sorted into endosomes and directed to various destinations. Rab GTPases, localized to distinct endosomes, regulate these processes by controlling membrane budding, vesicle formation, motility, tethering, and fusion through effector recruitment (11). It has been shown that HAdV-C2 and -C5 reach the cytosol in a Rab5-independent manner, while HAdV-C2_ts1 depends on Rab5 for endocytosis and transport to late endosomes and lysosomes. At the same time, sorting of HAdV-C2_ts1 to late endosomes was independent of Rab7 (12). To test if functional Rab proteins are involved in HAdV-D26 intracellular trafficking, we determined HAdV-D26 transduction efficiency in cells transfected with wild-type (WT) or dominant-negative (DN) form of Rab5, Rab7, Rab9, or Rab11. Dominant-negative mutants work in cells by competing with endogenous Rabs for binding to GEFs (Guanine Nucleotide Exchange Factor) and since they cannot interact with downstream target proteins, they form ‘dead-end’ complexes further preventing the activation of endogenous Rabs (21). Transduction efficiency of HAdV-D26 in cells transfected with DN form of Rab5, Rab7, or Rab9 measured by flow cytometry (Fig. 5A) and confocal microscopy (Fig. 5B) was partially decreased, while Rab11 DN did not influence HAdV-D26 transduction efficiency. The most pronounced effect was observed in cells expressing Rab9 DN, where HAdV-D26 transduction efficiency was decreased by approximately 40% compared to cells transfected with Rab9 WT. These results suggest that HAdV-D26 transduction efficiency partially depends on functional Rab5, Rab7, and Rab9 proteins.

**Figure 5.**
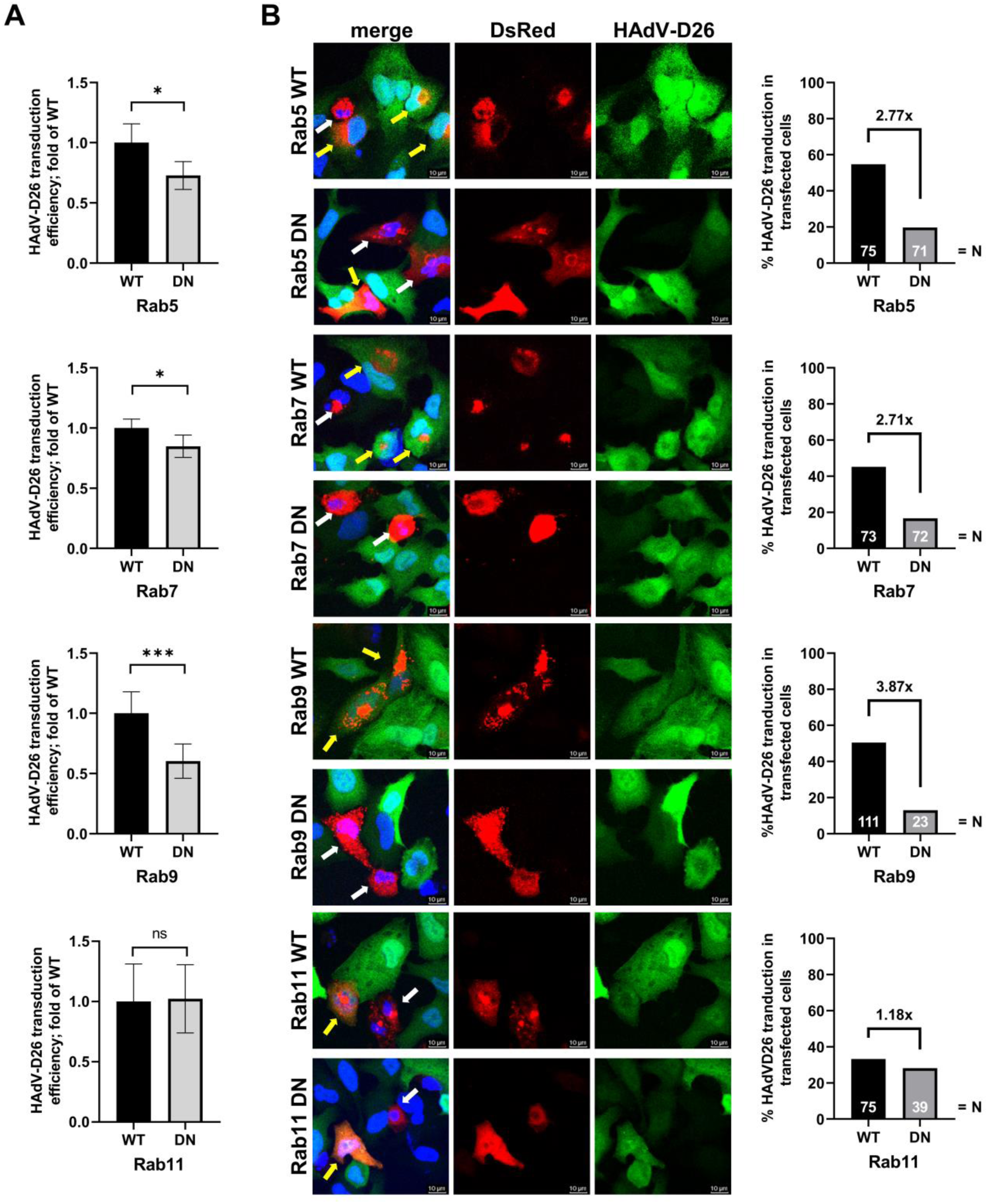
HAdV-D26 transduction efficiency partially depends on functional Rab5, Rab7 and Rab9 proteins. A549 cells were transfected with WT or DN Rab5, Rab7, Rab9 or Rab11 plasmids (48 h, 37°C), and subsequently incubated with HAdV-D26 for one hour, after which the medium was changed, and fresh medium was added. After 24 h, transduction efficiency was assessed by flow cytometry or confocal microscopy. A) HAdV-D26 transduction efficiency corresponding to the geometric mean of the fluorescence intensity of eGFP signal encoded by HAdV-D26 (MOI 10^4^ viral particles per cell) shown as a value relative to the corresponding WT sample. Data are presented as mean ± SD from three independent experiments in duplicates. B) Panels show representative cells from one experiment, where cells expressing a certain Rab protein (DsRed positive cells; red) were scored for HAdV-D26 infection (eGFP positive cells; green). Cell nuclei are shown in blue (DAPI). Yellow arrows indicate cells that are red/green positive (transfected and transduced cells), while white arrows indicate cells that are red positive (transfected cells) and green negative (non-infected cells). Scale bar, 10 μm. HAdV-D26 transduction efficiency (MOI 10^5^ vp/cell) is quantified as a percentage of the eGFP expressing cells transfected with Rab5, Rab7, Rab9 or Rab11 DN compared to eGFP expressing cells transfected with WT plasmids (right hand graphs). N represents the number of analyzed cells. *, P < 0.05; ***, P < 0.001.

### Rab9 knockdown decreases HAdV-D26 localization in the lysosome

To investigate further the role of Rab proteins in HAdV-D26 intracellular trafficking, we analyzed HAdV-D26 co-localization with EEA1 in cells transfected with siRNA specific for Rab5, Rab7, or Rab9 (Fig. 6A) (Rab5, Rab7 or Rab9 knockdown efficacy is shown in Fig. S2). The results show that knockdown of Rab5 protein in A549 cells increases the localization of HAdV-D26 in the early endosome. Namely, at 30 min p.i., 13% more HAdV-D26 localized to EEA1-positive endosomes in Rab5 knockdown cells compared to those transfected with scrambled siRNA. However, by 120 min p.i., HAdV-D26 localization in EEA1-positive endosomes was similar between Rab5 knockdown and control cells (si(−)). Rab7 knockdown in A549 cells did not affect HAdV-D26 localization in early endosomes at 30 min p.i., but during the following 90 min, it increased retention of HAdV-D26 in early endosomes compared to control cells.

**Figure 6.**
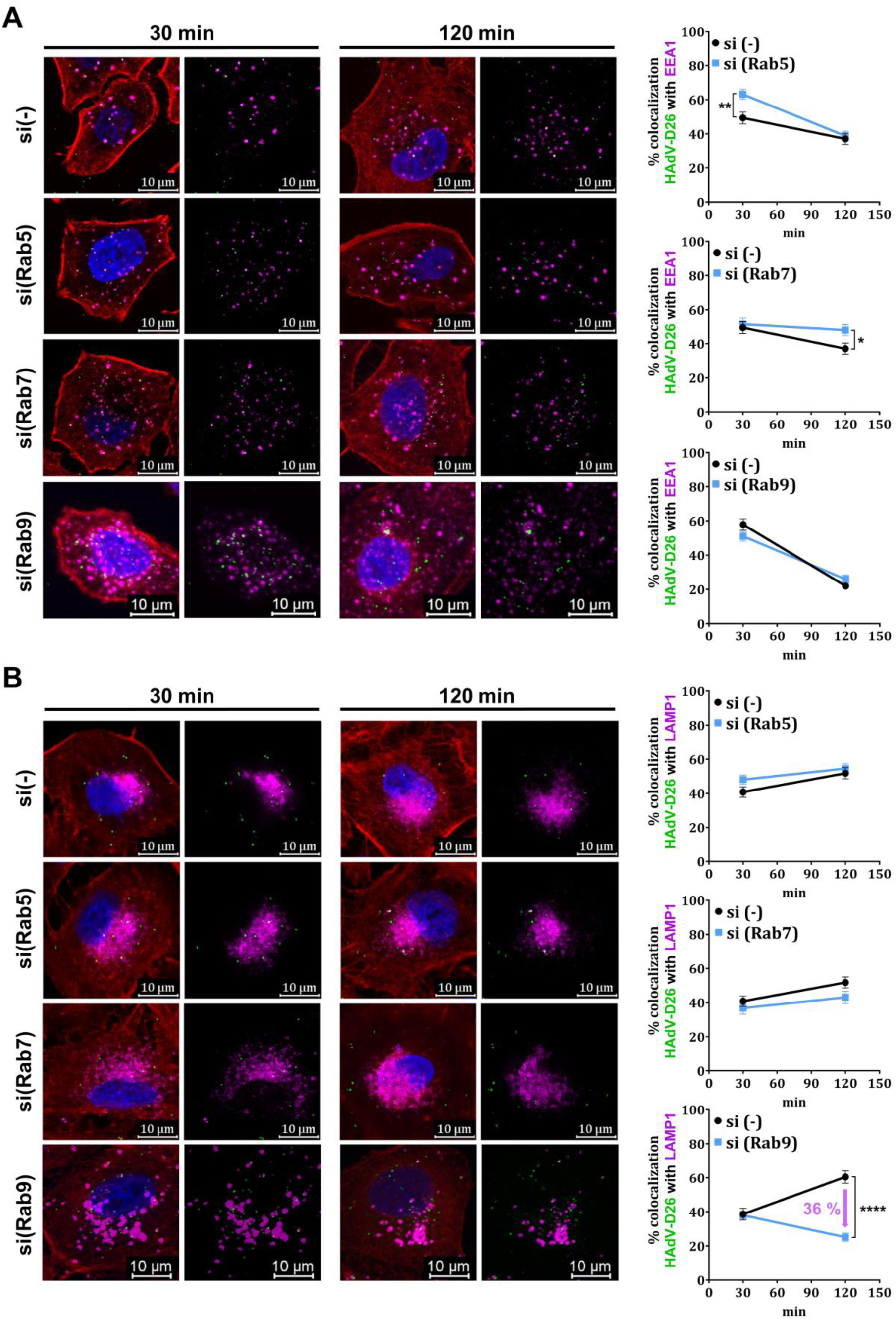
Rab5 and Rab7 knockdown increase HAdV-D26 localization within early endosomes, while Rab9 knockdown decreases HAdV-D26 localization within late endosomes/lysosomes. Cells were transfected with specific siRNA at the final concentration of 10 nM for Rab5 and Rab7, and 5 nM for Rab9, and 48 h later incubated with fluorescently labeled HAdV-D26 (green; MOI of 10^5^ vp/cell) for 45 min on ice to ensure uniform virus entry into the cell and subsequently transferred to 37°C for 10 min. Unbound viruses were washed, fresh medium was added, and cells were returned to 37°C for 30 min or 120 min, after which they were fixed and stained with antibody against (A) EEA1 (magenta) or (B) LAMP1 (magenta). F-actin is shown in red, and the nucleus in blue. Representative confocal images are shown. Scale bar = 10 μm. Graphs on the right show quantification of HAdV-D26 co-localization with EEA1 or LAMP1 at 30 and 120 min following knockdown of the indicated Rab protein. Colocalization was quantified by counting the number of virus particles overlapping with EEA1- or LAMP1-positive structures per cell. Data represent the mean ± SEM from at least 30 cells analyzed per condition. *, P < 0.05; **, P <0.01; ****, P < 0.0001.

At specific time points, increased co-localization of HAdV-D26 with EEA1 in Rab5 or Rab7 knockdown cells suggests that, under normal conditions, HAdV-D26 might be liberated from Rab5 and Rab7 positive early endosomes. Knockdown of Rab9 does not change the percentage of HAdV-D26 co-localization with EEA1 marker, indicating that Rab9 does not interfere with HAdV-D26 trafficking within early endosomes.

Next, we analyzed HAdV-D26 co-localization with LAMP1 in cells transfected with siRNA specific for Rab5, Rab7, or Rab9 (Fig. 6B). Knockdown of Rab5 protein in A549 cells modestly increases the localization of HAdV-D26 in the late endosome/lysosome at 30 min p.i., with 10% more virus localized in LAMP1-positive endosomes compared to cells transfected with scrambled siRNA. At 120 min p.i., HAdV-D26 localization in LAMP1-positive endosomes was similar between Rab5 knockdown and control cells. In contrast, Rab7 knockdown in A549 cells reduced the proportion of HAdV-D26 in late endosomes/lysosomes at 120 min p.i. from 52% to 42%. This decrease in localization in LAMP1 positive endosomes is even more obvious in cells with Rab9 knockdown, where localization of HAdV-D26 120 min p.i. decreased from 60% to 24%. Decreased co-localization of HAdV-D26 with LAMP1 in Rab7 or Rab9 knockdown cells suggests that Rab7 and Rab9 might promote HAdV-D26 delivery to late endosome/lysosome. Knockdown of Rab5 does not change the percentage of HAdV-D26 co-localization with LAMP1 marker, indicating that Rab5 does not interfere with HAdV-D26 trafficking within late endosomes/lysosomes.

### Rab9 knockdown increases HAdV-D26 transduction efficiency

Since we saw that functional Rab proteins are involved in HAdV-D26 transduction and that knockdown of Rab proteins influences HAdV-D26 transport through the endosomes, we wanted to investigate if depleting different Rab proteins would also play a role in the transduction efficiency of HAdV-D26. Thus, we downregulated Rab5, Rab7, or Rab9 by using specific siRNAs and measured the transduction efficiency of HAdV-D26. As shown in Fig. 7A, a decrease of approximately 30% in HAdV-D26 transduction efficiency was observed upon Rab5 downregulation, while downregulation of Rab7 did not influence HAdV-D26 transduction efficiency. On the other hand, downregulating Rab9 increased HAdV-D26 transduction efficiency by 62%. Increased adenovirus transduction efficiency can be a consequence of increased internalization or increased adenovirus genome delivery to the host nucleus (4). Therefore, we first investigated whether decreased expression of Rab9 influenced HAdV-D26 internalization. We quantified the number of internalized fluorescently labeled HAdV-D26 30 and 120 min p.i. and observed that downregulation of Rab9 increased HAdV-D26 internalization in average by more than 30% (Fig. 7B). Additionally, knowing that Rab9 knockdown increases HAdV-D26 escape from lysosomes, we also wanted to investigate if Rab9 knockdown would change the amount of HAdV-D26 genome imported into the host nucleus. For that purpose, we measured HAdV-D26 genome delivery to the nucleus in A549 cells transfected with scrambled or Rab9 siRNA and observed an increase in HAdV-D26 DNA nuclear import in cells with Rab9 knockdown (Fig. 7C).

**Figure 7.**
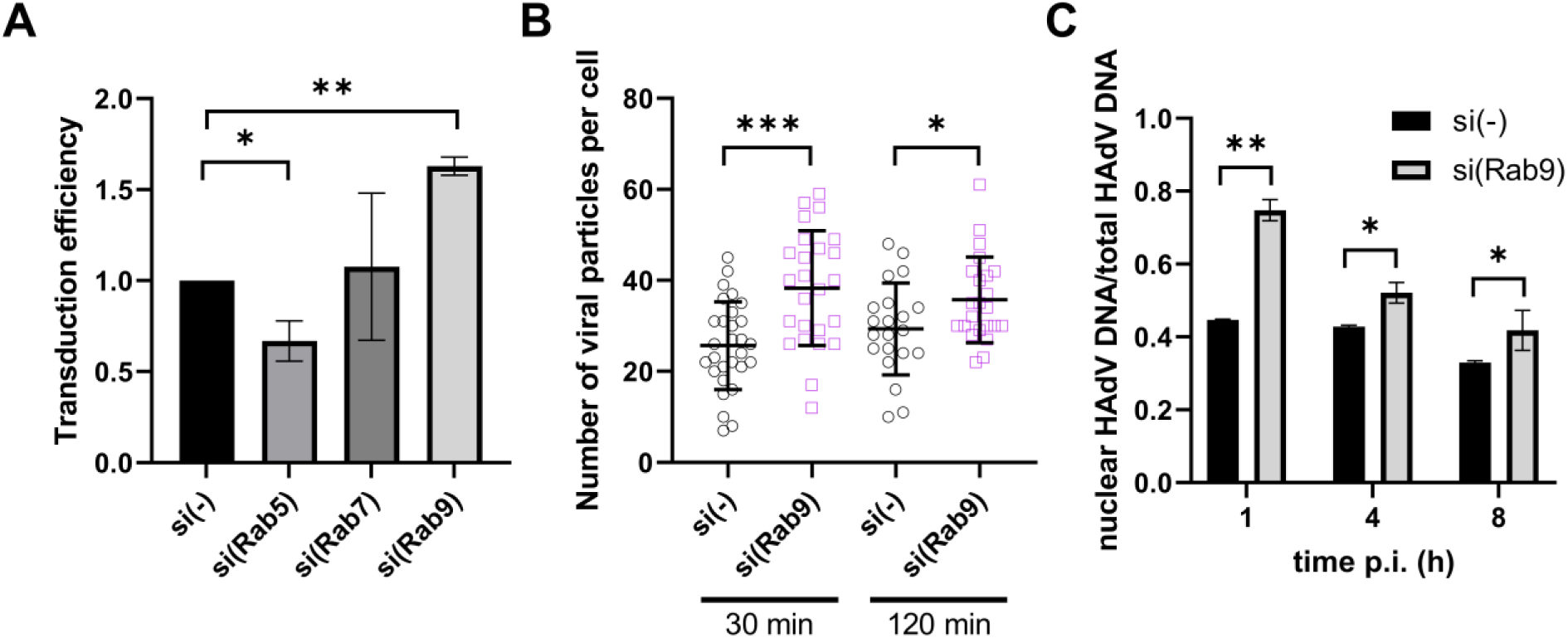
Knockdown of Rab9 increases internalization and transduction efficiency of HAdV-D26. A) Transduction efficiency of HAdV-D26 in A549 cells downregulated for Rab5, Rab7, or Rab9. Cells were transfected with specific siRNA at a final concentration of 10 nM for Rab5 and Rab7, and 5 nM for Rab9, and 48 h later infected with HAdV-D26 at an MOI of 10^4^ vp/cell. Transduction efficiency was measured as eGFP fluorescence 48 h p.i. The results are obtained by flow cytometry and presented as mean ± SD from three independent experiments normalized to control, i.e., cells transfected with scrambled siRNA. B) Internalization of HAdV-D26 in A549 cells downregulated for Rab9. Cells were transfected with specific Rab9 siRNA, 5 nM, and 48 h later infected with fluorescently labeled HAdV-D26, MOI 10^5^ vp/cell, for 45 min on ice to ensure uniform virus entry into the cell and subsequently transferred to 37°C for 10 min. Unbound viruses were washed, fresh medium was added, and cells were returned to 37°C for 30 min or 120 min, after which they were fixed and imaged by confocal microscopy. Quantification is presented as the number of viral particles per cell with mean ± SD. C) HAdV-D26 genome delivery into the host nucleus in Rab9 downregulated cells. Cells were transfected with scrambled siRNA or Rab9 siRNA at a final concentration of 5 nM, and 48 h later infected with HAdV-D26, MOI 10^4^ vp/cell for 45 min on ice to ensure uniform virus entry into the cell. Five min after incubation on 37°C unbound viruses were removed, and fresh medium was added, followed by incubation at 37°C. Cells were harvested at defined time intervals. DNA was extracted from the total cells or isolated nuclei. Viral and cellular DNA were quantified by qPCR using primers for the CMV promoter present in the viral DNA or the cellular GAPDH gene. Results are presented as a ratio of viral DNA measured in the nuclear fraction to the total fraction, normalized to the corresponding si(−) value. Data are presented as mean values ± SD from three experiments. Subsequently, the same experimental setup was performed as described in Figure 2A. *, P < 0.05; **, P < 0.01, ***, P < 0.001.

## Discussion

HAdV-D26 was first isolated in 1956 from an anal specimen of a 9-month-old child, potentially associating it with enteric infections (22). In contrast to HAdV-C5, HAdV-D26 has a low seroprevalence in humans, which has promoted its development as a vaccine vector (3). The functionality of HAdV-D26–based vectors is strongly influenced by receptor usage and intracellular trafficking. HAdV-D26 can utilize multiple entry receptors with receptor preference depending on the host cell type (5, 23, 24). However, none of these receptors appears to bind HAdV-D26 with high affinity, consistent with the higher MOI required to achieve infectivity comparable to HAdV-C5 or HAdV-B35 (5). Although adenoviruses typically enter cells via clathrin-mediated endocytosis (25, 26), HAdV-D26 preferentially uses a caveolin-mediated pathway in cells expressing high levels of αvβ3 integrin (6). In contrast, in cells with low αvβ3 integrin expression, inhibition of clathrin-mediated endocytosis impairs viral internalization, indicating that entry pathways are receptor dependent (6). Despite its importance as a vaccine vector, HAdV-D26 intracellular trafficking remains poorly characterized, with only one study reporting late endosomal trafficking (7). Elucidating these trafficking routes is critical for optimizing vector performance. Here, we investigated the post-internalization trafficking of HAdV-D26 in A549 cells, focusing on the role of endosomal compartments in viral trafficking and transduction efficiency.

Under *in vitro* conditions, HAdV-C2 and HAdV-C5 rapidly escape endosomes and reach the nucleus within 60 min post-infection (27). In contrast, HAdV-D26 reaches the nucleus considerably more slowly. Fluorescence imaging revealed that at 60 min post-infection, HAdV-D26 remains dispersed throughout the cytoplasm, and even at 240 min shows no pronounced nuclear accumulation, with particles instead clustering asymmetrically in the perinuclear region. A similar trafficking pattern has been reported for HAdV-B7, and chimeric Ad5f7 virions (chimeric vector composed of HAdV-C5 capsid and HAdV-B7 fiber), indicating altered endosomal escape (28), which may likewise apply to HAdV-D26. To map the intracellular route of HAdV-D26, we imaged HAdV-D26 intracellular trafficking by TEM during the first 60 min post-infection and detected viral particles at the plasma membrane, within vesicular structures, and free in the cytosol. Notably, nearly 40% of viral particles accumulated in degradative compartments, indicating prolonged endosomal retention, contrary to HAdV-C5, which escapes endosomes as early as 20 min after uptake (17).

Because TEM revealed substantial accumulation of HAdV-D26 in degradative compartments, likely lysosomes, we examined its endosomal trafficking in more detail by assessing co-localization with early (EEA1) and late endosome/lysosome (LAMP1) markers during the first 240 min post-infection. Within the first 120 min, ∼80% of HAdV-D26 particles remained in early or late endosomes/lysosomes, indicating limited endosomal escape. Although HAdV-D26 and HAdV-C5 both initially localize to early endosomes, co-infection experiments showed that they do not share the same endosomal pathway, as HAdV-C5 did not enhance HAdV-D26 transduction. Notably, over 40% of HAdV-D26 persisted in LAMP1- and LysoTracker-positive compartments for up to 240 min, suggesting impaired endosomal escape or trafficking into lysosomal compartments. Our imaging data are in line with a previously reported study proposing that HAdV-D26 accumulates in late endosomes 2 to 8 h post-infection (7). The authors show co-localization of HAdV-D26 with EEA1 and LAMP1 in A549 cells, however, they observed a much lower co-localization percentage of HAdV-D26 than we did. Prolonged stay within lysosomes was reported also for HAdV-C2_ts1, HAdV-B7, and Ad5/35L (chimeric vector composed of HAdV-C5 capsid and HAdV-B35 fiber knob). HAdV-C2_ts1 and Ad5/35L mediate significantly less efficient gene transfer compared to that of HAdV-C5. While HAdV-C2_ts1 fails to package the viral protease L3/p23, and therefore does not efficiently escape from endosomes (12), Ad5/35L used late endosomal/lysosomal compartments to achieve localization to the nucleus. However, a significant portion of Ad5/35L particles appeared to be recycled back to the cell surface (14). On the other hand, HAdV-B7 residence in late endosomes and lysosomes seems to be the native trafficking pathway, causing no loss of viral genome in low-pH compartments (29). Failure to escape the endosomes was reported for HAdV-C5 in murine alveolar macrophages, an effect attributed to GM-CSF via PU.1, which limits productive infection (30). Based on our results, we can conclude that 20% HAdV-D26 is liberated from any endosome, while more than 40% is retained in the late endosome/lysosome. We hypothesize that the remaining 40% is sorted to an unknown endosome from where it is either released or remains retained. A similar phenomenon was observed for HAdV-C2 in A549 cells, where endosomes with unreleased HAdV-C2 bypass normal early-to-late endosome maturation and either enter an unknown non-permissive compartment or recycle back to the plasma membrane (31). Since Rab11 DN had no influence on HAdV-D26 transduction efficiency and HAdV-D26 did not exhibit significant movement away from the nucleus, we conclude that HAdV-D26 does not follow the path of the recycling endosomes.

Lysosomes are the primary degradative compartments of eukaryotic cells, breaking down material internalized by endocytosis (32). HAdV-D26 that reaches lysosomes could be degraded and fail to reach the nucleus. We found that inhibiting lysosomal acidification, but not transport from early to late endosomes/lysosomes, modestly increased HAdV-D26 transduction. Similar effects were reported for HAdV-C5 and HAdV-B3, though the increase was much greater for those viruses (17). This suggests that low endosomal/lysosomal pH is not required for HAdV-D26 transduction. The modest increase in HAdV-D26 transduction efficiency in cells treated with bafilomycin A1 implies that HAdV-D26 entrapped in the lysosomes is lost in terms of transduction, presumably due to degradation. Overall, our results indicate that the transition from early endosome to lysosome is not a prerequisite for efficient transduction with HAdV-D26. Importantly, lysosomal residence may benefit HAdV-D26 vaccine vectors by promoting antigen processing for MHC class II presentation (33) and engaging innate immune sensors (34), providing a self-adjuvanting effect that could enhance adaptive immunity.

Multiple entry pathways allow viruses to adapt to different cell types, evade host defenses, and maximize infection. Some pathways lead to productive nuclear entry, while others result in degradation. HAdV-B7-based vectors transfer genes as efficiently as HAdV-C5 vectors, despite trafficking through late endosomes/lysosomes, showing that species B adenoviruses can deliver their genome effectively via distinct pathways (14). Contrary to both HAdV-C5 and HAdV-B7, here we demonstrated that HAdV-D26 is rather inefficient in nuclear delivery of its genome. Namely, the highest amount of HAdV-D26 DNA imported into the host nucleus was observed 1 h post-infection, while in the later infection time points, the DNA amount slightly decreased, indicating an issue with nuclear import. Adenoviral DNA nuclear import requires capsid disassembly and interactions between viral and host factors, including hexon, protein V/VII, Mind bomb 1, Nup214, kinesin, and others (15, 35, 36). Disruption of these interactions can block nuclear entry, leaving viral genomes in the cytoplasm or subject to degradation. Since we did not measure any decrease in the total amount of HAdV-D26 DNA that entered the cell over the course of 8 hours, we assume that HAdV-D26 DNA is not degraded but rather stays in the cytoplasm. We saw that only approximately 34% of rapidly moving HAdV-D26 reaches the nucleus which could also account for low nuclear import.

Rab proteins define endosomal identity and regulate vesicle budding, transport, tethering, and fusion through effector recruitment. Their spatially and temporally regulated, reversible assembly enables dynamic changes in membrane composition and organelle fate (37). In A549 cells, a significant portion of HAdV-D26 traffics to late endosomes/lysosomes, a process independent of Rab5 and Rab7 but dependent on Rab9, as Rab9 knockdown reduced HAdV-D26 co-localization with LAMP1 by over 35%. Rab9 knockdown also increased HAdV-D26 transduction efficiency, internalization, and genome import into the host nucleus, suggesting that Rab9 limits HAdV-D26 escape from lysosomes. Notably, transduction results with DN Rab9 differed from those observed after Rab9 downregulation. The divergent effects observed with Rab9 knockdown and expression of DN Rab9 likely reflect the fundamentally different nature of these perturbations. siRNA produces a specific reduction in Rab9 levels and may promote accelerated exit from late endosomes, favoring HAdV-D26 endosomal escape, or may shift HAdV-D26 to an alternative route. On the other hand, DN Rab9 sequesters upstream GEFs and exerts broader inhibitory effects on endosomal maturation and related Rab pathways, leading to a more global disruption of late endosomal trafficking. Such pleiotropic effects can impair the membrane remodeling steps required for HAdV-D26 penetration, resulting in reduced infection.

Rab9 normally facilitates late endosome-to-trans-Golgi trafficking and lysosomal degradation of cargo (38), while Rab9 depletion alters endosome maturation by enlarging late endosomes and disrupting cargo retention (39). Therefore, Rab9 knockdown could increase HAdV-D26 infection by impairing the Rab9-dependent lysosomal degradation pathway, reducing lysosomal delivery of HAdV-D26 particles, and enabling their accumulation in enlarged endosomes, promoting cytoplasmic escape. For example, Rab9 has been shown to mediate the trafficking of hepatitis B virus envelope proteins to lysosomes for degradation via interaction with autophagy receptor NDP52. In the same study, depletion of Rab9 resulted in increased hepatitis B virus DNA species in the cells, indicating that downregulation of the Rab9-dependent lysosomal degradation pathway promotes HBV replication (40). Conversely, Rab9 knockdown blocks HPV entry by disrupting retromer-mediated endosome-to-Golgi transport, causing viral accumulation in endosomes (41). In the same study, authors report that the GDP-bound form of Rab9 actually promotes HPV entry, whereas GTP-bound Rab9a inhibits it, indicating that Rab9 GTPase nucleotide state has non-intuitive effects on infection and can behave non-canonically when it comes to viral cell entry. To the best of our knowledge, Rab9 has not been described so far as a limiting factor for adenovirus transduction efficiency.

From the literature we know that Rab9 can act as an oncogenic driver and therapeutic target in cancers such as breast cancer (42), melanoma (43), and gastric cancer (44) where Rab9 knockdown inhibited proliferation, migration, or invasion of cancer cells. Since Rab9 downregulation enhances HAdV-D26 transduction, HAdV-D26-based vectors could serve as combination or second-line therapies alongside Rab9 downregulation, particularly in oncolytic virotherapy, where transient Rab9 silencing would boost vector’s efficacy.

Providing more details regarding the possible degradation of HAdV-D26 within the lysosomes was beyond the scope of this study, however, our data raises the question why HAdV-D26 would end up in the lysosomes in the first place. It is possible that HAdV-D26 can use the receptor which would sort it to the lysosomes, however none of the molecules that have been so far proposed to be involved in HAdV-D26 cell entry do not actively direct their ligands to lysosomes. One of the molecules shown to direct proteins to the lysosome for degradation is the mannose 6-phosphate receptor (M6PR) (45, 46). Herpes viruses 1 and 2, varicella zoster virus, and human immunodeficiency virus 1 can exploit the M6PRs located on the cell surface as an entry receptor or co-receptor for invading host cells (47). Knowing that Rab9 has been reported to be involved in sorting M6PR to late endosomes (38), one can wonder if HAdV-D26 could maybe interact with M6PR speculating that this interaction would route it towards late endosomes/lysosomes in Rab9 dependent manner. Once Rab9 is downregulated, HAdV-D26 is not sorted to late endosomes/lysosomes but rather uses an alternative pathway, resulting in better transduction efficiency, as seen in our results. However, this alternative pathway still needs to be described. HAdV-D26 was originally isolated from stool, although it is not classified as a classical enteric adenovirus. Its retention within late endosomes may nevertheless indicate an increased tolerance for acidic intracellular environments, potentially reflecting structural features compatible with gastrointestinal exposure.

Our results extend current knowledge of adenovirus trafficking by providing new insights into post-entry HAdV-D26 dynamics and the role of Rab proteins in HAdV-D26 transduction efficiency. We highlight distinctive features of HAdV-D26, including prolonged lysosomal residence and limited genome delivery to the nucleus. Although Rab9 modulation affects viral internalization and transduction (Fig. 8), the precise molecular mechanisms remain unclear. A limitation of our study is that the primary HAdV-D26 receptor is still unknown, so epithelial cell models may not fully reflect *in vivo* conditions. Nevertheless, these findings offer valuable insights into virus-host interactions that can guide the design of safer and more effective adenoviral vectors.

**Figure 8.**
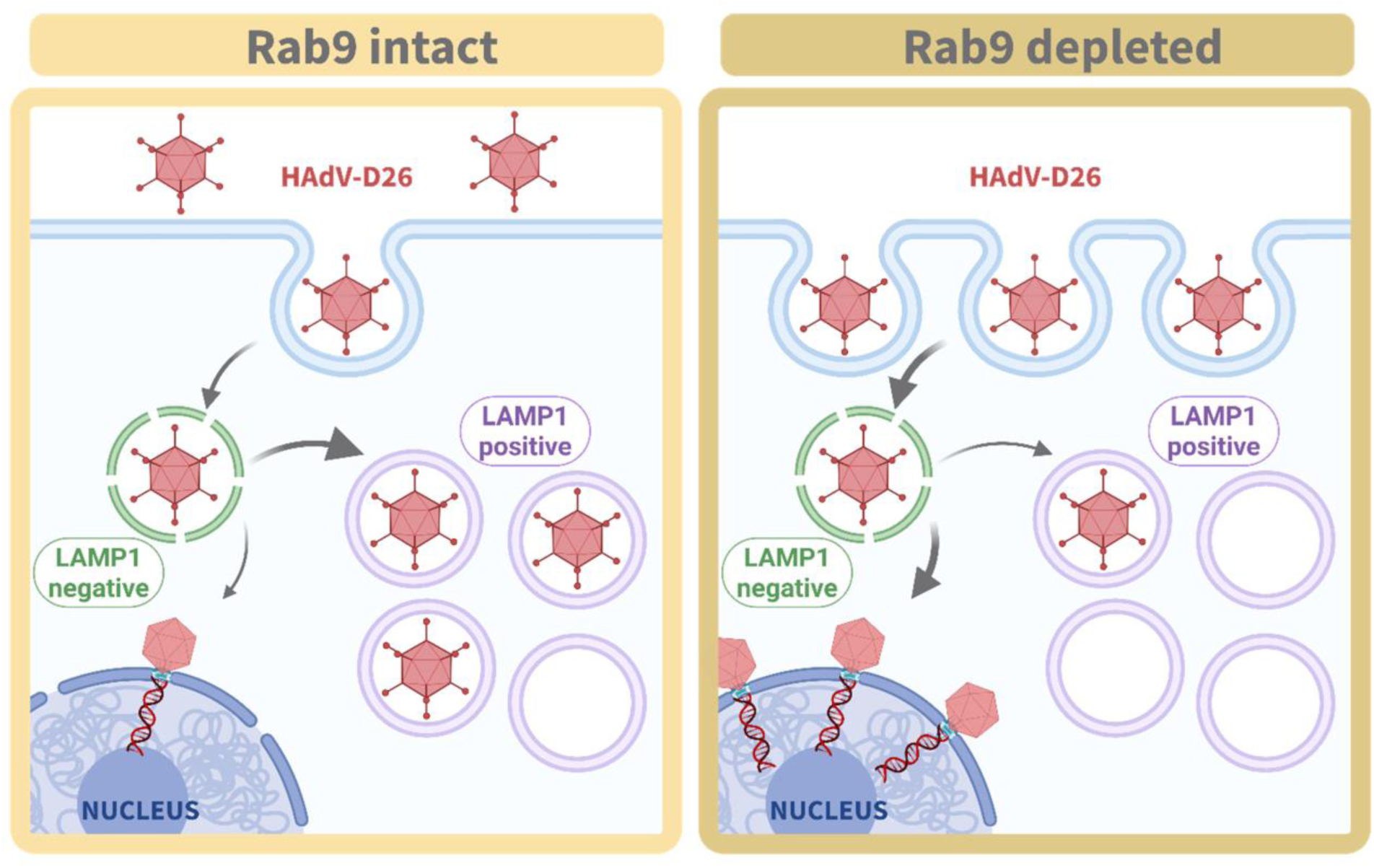
Schematic presentation of how Rab9 influences HAdV-D26 intracellular trafficking and transduction efficiency. Decreased expression of Rab9 allows for increased HAdV-D26 internalization but also increased liberation from currently unidentified endosomal compartment. Subsequently, HAdV-D26 retention in late endosome/lysosome (LAMP1-positive) is decreased, which could decrease potential HAdV-D26 degradation and promote better transduction efficiency. This illustration was created in BioRender (https://www.biorender.com/).

## Materials and methods

### Cells, viruses, and antibodies

HEK293 (human embryonic kidney; ATCC CRL-1573) and A549 (human lung carcinoma; ATCC CCL-185) cells were obtained from the ATCC Cell Biology Collection and were cultured in antibiotic-free Dulbecco’s Modified Eagle’s Medium- high glucose (DMEM) supplemented with 10% (vol/vol) Fetal Bovine Serum (FBS) if not stated differently. Osteosarcoma U2OS cell line (female) stably expressing mCherry-α-tubulin was a kind gift from Marin Barišić (Danish Cancer Society Research Center, Copenhagen, Denmark) and Helder Maiato (Institute for Molecular Cell Biology, University of Porto, Portugal). U2OS cells were maintained in DMEM supplemented with 10% (vol/vol) FBS and 50 µg/mL geneticin. All cells were grown at 37°C in a humidified incubator with a 5% CO_2_ atmosphere. Replication-incompetent recombinant adenoviral vectors based on adenovirus types 2, 5 or 26 were previously constructed (48, 49). Viruses were propagated on HEK293 cells and purified by CsCl gradients. They carried an enhanced green fluorescent protein (eGFP) gene driven by the cytomegalovirus (CMV) promoter as a reporter gene. Fluorescent labeling of adenoviruses was previously described (5). Primary antibodies used for immunofluorescence and co-localization analyses, or western blot were as follows: anti-EEA1 (#2411, Cell Signaling, SAD), anti-LAMP1 (ab24170, Abcam, UK), anti-Rab5 (#3547, Cell Signaling, SAD), anti-Rab7 (#9367, Cell Signaling, SAD), anti-Rab9A (#5118, Cell Signaling, SAD), anti-Alexa-Fluor-488 recombinant F(ab)-rabbit monoclonal antibody (15H9L93, Invitrogen).

### Transduction efficiency assay

Transduction efficiency was measured either by flow cytometry as previously described (6) or by spectrophotometer. For assaying by spectrophotometer, cells were seeded in 96-well plates at a density of 4×10^3^ cells per well, and 48 h later incubated with HAdV vectors at MOI of 10^4^ vp/cell. After 1 h at 37°C in a humidified incubator with a 5% CO_2_ atmosphere the medium was removed and replaced with of antibiotic-free DMEM supplemented with 10% (vol/vol) FBS and cultured for additional 24 h. Cells were then washed twice with PBS and eGFP expression was measured by spectrophotometer at 515 nm. Subsequently, after washing, cellular DNA was stained with Hoechst 33342 (5 µM) and the fluorescence intensity was measured at 455 nm. The results were calculated as the ratio of eGFP/Hoechst values.

For measuring transduction efficiency after interfering with endosome pH cells were pretreated with following inhibitors (30 min, 37°C): bafilomycin A1 (10 nM), chloroquine (50 µM), NH_4_Cl (5 mM) or EGA (15 µM) and subsequently incubated with HAdV-D26 for one hour (MOI 10^4^ vp/cell) in medium with the specific inhibitor, after which the medium was changed and fresh medium was added. After 24 h of incubation, the fluorescence intensity of eGFP was measured and subsequently the fluorescence intensity of cellular DNA labeled by Hoechst.

For studying role of Rab proteins in HAdV-D26 transduction efficiency cells were transfected with DN or WT form of Rab5, Rab7, Rab9 or Rab11 purchased from Addgene: DsRed-Rab5 WT, #13050; DsRed-Rab5 DN, #13051; DsRed-Rab7 WT, #12661; DsRed-Rab7 DN, #12662; DsRed-Rab9 WT, #12677; DsRed-Rab9 DN, #12676; DsRed-Rab11 WT, #12679; DsRed-Rab11 DN, #12680. Plasmids were transfected into A549 cells using Lipofectamine (Thermo Fisher Scientific) according to the manufacturer’s protocol, and 48 h after cells were transduced with HAdV-D26.

### SLO penetration assay

This assay has been previously described (17). Briefly, A549 cells were seeded on Alcian blue-coated glass coverslips in 24-well dishes (40,000 cells/well) and grown for 2 days. Alexa-488-labeled viruses were bound to cells at 0°C for 60 min. The unbound virus was washed away and the cells were placed into a 37°C water bath for 45 min. After virus internalization, cells were placed on ice, washed twice with streptolysin O (SLO) binding buffer and processed for SLO-mediated perforation of the plasma membrane. Cells were washed twice with SLO binding buffer and incubated with SLO solution (6.2 µg/mL) for 10 min on ice. Unbound SLO was removed by two washes with SLO binding buffer. Cells were then incubated in SLO binding buffer for 5 min at 37°C in a water bath to induce pore formation and immediately returned to ice. After being washed with SLO internalization buffer cells were incubated with anti-Alexa-488 antibody for 1 h at 0°C. Next, cells were and fixed with 3% paraformaldehyde, permeabilized with 0.5% Triton X-100 in PBS for 5 min and washed once with PBS. Blocking was performed with 10% goat serum in PBS for 5 min at room temperature. Then, cells were incubated with anti-rabbit Alexa594 antibody and DAPI for 30 min at room temperature, followed by three 4-min washes in PBS. A post-fixation step was performed with 3% PFA in PBS for 12 min at room temperature. Cells were washed once with PBS, quenched with 25 mM ammonium chloride in PBS for 10 min, washed once more with PBS, and mounted for imaging.

### siRNA experiments

To downregulate specific molecules, we used following predesigned MISSION® esiRNA: Rab5 (EHU053901), Rab7 (EHU053221), Rab9 (EMU171281), universal negative control #1 (SIC001), all from Sigma-Aldrich. Cells were transfected at a confluence of 30 to 50% using Lipofectamine RNAiMax reagent (Invitrogen) according to the manufacturer’s protocol. Final esiRNA concentrations were 10 nM for Rab5 and Rab7, and 5 nM for Rab9. The efficiency of silencing was verified 48 h after transfection by western blot. Briefly, cells transfected with specific or negative control siRNA were lysed with Laemmli buffer heated to 95°C and sonicated. Proteins were separated by SDS-PAGE and transferred to nitrocellulose membrane. After blocking, the membrane was probed with appropriate primary antibodies, followed by incubation with appropriate horseradish peroxidase-conjugated IgG secondary antibody. Detection was performed with Pierce ECL Western Blotting Substrate using ChemiDoc Imaging System. Densitometry was performed with ImageJ software. Proteins were normalized to the total proteins stained with amido black. The results are presented as the relative expression of proteins compared to cells transfected with control siRNA.

### Transmission electron microscopy

A549 cells grown on grids were infected with HAdV-D26 at an MOI of 2×10^5^ vp/cell, incubated on ice for 45 min, and then shifted to 37°C for 10 min. After removal of unbound viruses by washing with fresh medium, cells were incubated for 1 h at 37 °C, after which they were washed twice with PBS and fixed with a mixture of 2% glutaraldehyde and 1% tannic acid in 0.4 M HEPES buffer (pH 7.2). Dehydration was performed using dry ethanol, and heavy metal staining and embedding in EPON resin were carried out on the monolayer according to the protocol described in (50). Serial sections 70 nm thick were collected from the first 2 µm of the basal cell surface (4 sections per grid, for a total of 9 grids). Sections were stained with uranyl acetate and lead citrate for image acquisition, which was performed using a JEOL JEM 1400 transmission electron microscope operated at 120 kV. The number of viral particles present in different cellular compartments was quantified in 56 cells, each one presenting an area greater than 50 µm². The distribution of viral particles among the different compartments was normalized to the total number of quantified particles (262 viral particles).

### Confocal microscopy

Cells (2×10^4^ per coverslip) were seeded on coverslips in 24-well plates. Two days later, fluorescently labeled adenoviruses were added to the cells (MOI 5×10^4^ vp/cell) and incubated on ice for 45 min to allow binding. Subsequently, the cells were transferred to 37°C for the indicated time. The cells were fixed with 2% paraformaldehyde (PFA) in PBS for 12 min at room temperature. Nuclei were labeled with DAPI (4=,6-diamidino-2-phenylindole). Coverslips were slide mounted using Fluoromount G (Southern Biotech). Confocal laser scanning microscopy analyses were performed using a Leica TCS SP8 X inverted confocal microscope (Leica Microsystems, Wetzlar, Germany) with 63x/1.40 oil-immersion objective. The images showing intracellular trafficking of Alexa Fluor 488-labeled HAdVs are maximum projections of at least 7 confocal stacks and were processed with the Leica Application Suite X (LAS X) software platform, Adobe Photoshop CC software (Adobe Systems), and ImageJ.

### Live cell microscopy

Day before live-cell microscopy, 6×10^4^ U2OS-mCherry-tub cells were seeded per chamber on 4-chamber glass-bottom dishes (#1.5 glass thickness, iBL). Viral infection was performed on ice with 5×10^4^ MOI and the cells were kept on ice until live cell imaging. The microscopy was performed on Expert Line easy3D STED microscope system (Abberior Instruments) with a 100 x/1.4NA UPLSAPO100x oil objective (Olympus, Tokio, Japan) and avalanche photodiode detector. During imaging, the cells were kept at 37°C and a 5% CO_2_ atmosphere within a heating chamber (Okolab). Images were acquired every ten seconds using Imspector software. Imaging was performed within an hour after infection. Image analysis was performed in Fiji/ImageJ (National Institutes of Health). Quantification and data analysis were performed in GraphPad Prism (GraphPad Software). Tracking of viral particles was done manually by using the Multi-point tool in Fiji. Upon inspection of viral particle movement within the cells, viral particles which could be followed for at least 20 s (three time- frames) were taken into account. Only fast movements towards or from the nucleus were tracked. For evaluation of virus redistribution, virus particles were counted as reaching the nucleus if they were ≤ 1 µm from the nuclear surface.

For co-localization of HAdV-D26 with LysoTracker in live A549 cells, 2×10^4^ cell were seeded on coverslips in 24-well plates. Two days later, fluorescently labeled adenoviruses were added to the cells (MOI 5×10^4^ vp/cell) and incubated on ice for 45 min to allow binding, then 10 min at 37°C, followed by washing with fresh medium to remove unbound viruses, and then incubated at 37°C for the indicated times. In the last 30 min of incubation, LysoTracker Deep Red solution (Invitrogen) was added to the medium to final concentration of 50 nM. After incubation, cells were washed with fresh medium and immediately observed by a Leica TCS SP8 X inverted confocal microscope (Leica Microsystems, Wetzlar, Germany) as stated before. Reflection Interference Contrast Microscopy (RICM) was used to determine cell shape. The colocalization analysis was performed using digital images processed with a colocalization plug-in in Fiji. Manders’ coefficient was used to quantify pixel-wise co-localization between HAdV-D26 and LysoTracker signals. Specifically, the coefficient represents the proportion of the HAdV-D26 signal overlapping with the LysoTracker signal, indicating the extent of viral localization within lysosomal compartments.

### Adenovirus genome delivery to the host nucleus

A549 cells (2×10^4^ per well) were seeded in 6-well plates, and 24 hours after seeding, viral infection was performed on ice with 1×10^4^ MOI in DMEM supplemented with 0.2% (vol/vol) FBS, and incubated on ice for 45 min to synchronize viral entry. After 5 min at 37°C in a humidified incubator with a 5% CO_2_ atmosphere, the medium was removed and replaced with antibiotic-free DMEM supplemented with 10% (vol/vol) FBS. The cells were then kept at 37 °C for defined time points (0, 1, 4, and 8 h).

To monitor viral DNA delivery dynamics in A549 cells after Rab9 silencing, Rab9 knockdown was performed using DharmaFECT 4 on the same day as cell seeding, according to the manufacturer’s protocol. After 48 hours of incubation, viral infection was performed as mentioned above. At the defined time points, cells were washed with PBS, trypsinized and after washing with PBS divided into two fractions. The cell pellet from first fraction was used for total DNA isolation, while the second cell pellet was used for nuclear fraction isolation. For nuclear fraction isolation, 100 µL of hypotonic solution (10 nM HEPES (pH 7.5), 10 mM NaCl, 1 mM DTT, 1 mM EDTA, 2 mM MgCl_2_x6H_2_0) was added to the pellet and incubated for 5 min at room temperature, followed by the addition of 100 µL of 0.05% NP-40 detergent for 10 min. The mixture was vortexed and centrifuged for 2 min at 13,400 × g. The resulting pellet was washed four times in PBS. The final slightly translucent pellet represented the nuclear fraction. Total cellular and viral DNA was extracted using the QIAamp DNA Blood Mini Kit (Qiagen) according to the manufacturer’s instructions. DNA concentration was measured spectrophotometrically at 260 nm, and purity was assessed at 280 nm. The isolated DNA served as a template for qPCR using primers targeting the CMV promoter (forward primer: 5’-TGGGCGGTAGGCGTGTA-’3; reverse primer: 5’-CGATCTGACGGTTCACTAAACG-3’) or the cellular GAPDH gene (forward primer: 5’-AGAACATCATCCCTGCCTCTACTG-3’; reverse primer: 5’ TGTCGCTGTTGAAGTCAGAGGAGA-3’).

### Statistical analysis

All experiments were performed in at least two biological replicas in duplicate or triplicate. All analyses and graphs were created in GraphPad Prism (GraphPad Software Inc., USA). Data were analysed by unpaired Student’s t test or Mann-Whitney U test and expressed as mean ± SD or SEM. ns, not significant; *, P < 0.05; **, P <0.01; ***, P < 0.001; ****, P < 0.0001.

## Funding

D.M. acknowledges funding from Croatia Science Foundation Research Projects (Grant IP-2019-04-6048 and IP-2025-02-3470 to D.M.); C.S.M. acknowledges funding from the Spanish State Research Agency, with co-funding from the European Regional Development Fund (PID2022-136456NB-I00/AEI/10.13039/501100011033). The C.S.M. group is a member of the Spanish Adenovirus Network (RED2022-134221-T/AEI/10.13039/501100011033). CNB-CSIC is an AEI Severo Ochoa Excellence Center (CEX2023-001386-S/AEI/10.13039/501100011033). Support from the European Commission Marie Skłodowska-Curie Actions (grant agreement 101129778, project INVECTA) to D.M. and C.S.M. is also acknowledged.

## Acknowledgements

We thank the CNB-CSIC Electron Microscopy facility for the excellent technical support, Urs Greber and Maarit Suomalainen for granting access to laboratory facilities and resources, and their valuable scientific input during the execution of the SLO experiment, Marin Barišić and Helder Maiato for kindly providing U2OS cell line stably expressing mCherry-α-tubulin, Marina Šutalo for her technical assistance and Lucija Horvat for confocal microscopy support.

## Author contributions

Conceptualization: Isabela Drašković, Davor Nestić, Dragomira Majhen

Data curation: Dragomira Majhen

Formal analysis: Isabela Drašković, Davor Nestić, Lucija Lulić-Horvat, Gabriela N. Condezo Dragomira Majhen

Funding acquisition: Dragomira Majhen

Investigation: Isabela Drašković, Davor Nestić, Lucija Lulić-Horvat, Jelena Martinčić, Mario Stojanović, Gabriela N. Condezo, Klara Kašnar

Methodology: Isabela Drašković, Davor Nestić, Lucija Lulić-Horvat, Jelena Martinčić, Mario Stojanović, Gabriela N. Condezo, Klara Kašnar, Carmen San Martín, Jerome Custers, Dragomira Majhen

Resources: Carmen San Martín, Jerome Custers, Dragomira Majhen Supervision: Dragomira Majhen

Visualization: Isabela Drašković, Davor Nestić, Lucija Lulić-Horvat, Jelena Martinčić, GabrielaN. Condezo

Writing – original draft: Isabela Drašković, Davor Nestić, Dragomira Majhen

Writing – review & editing: Isabela Drašković, Davor Nestić, Jelena Martinčić, Mario Stojanović, Gabriela N. Condezo, Carmen San Martín, Dragomira Majhen

## Supplementary figures

**Figure S1.**
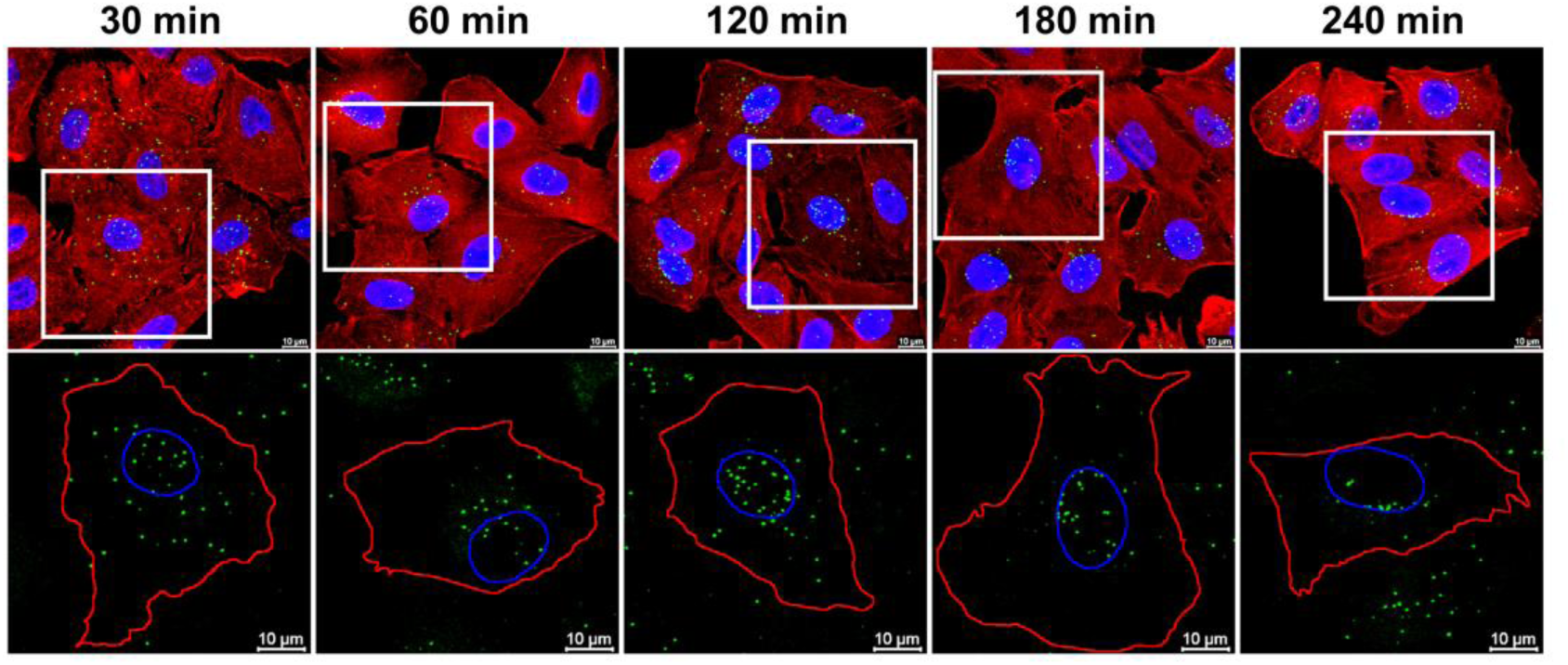
Intracellular trafficking of HAdV-C5 in epithelial cells. Cells were incubated on ice with fluorescently labeled HAdV-C5 (green; MOI 5×10^3^ vp/cell) for 45 min, then 10 min at 37°C, followed by washing with fresh medium to remove unbound viruses. After additional incubation at 37°C for the indicated times, cells were fixed with 2% PFA/PBS and stained with phalloidin-AF555 (red) and DAPI (blue). Representative confocal images are shown (scale bar = 10 μm). The lower panel represents increased selection from the upper panel, with marked edges of the nucleus (blue, based on DAPI) and cell (red, based on phalloidin).

**Figure S2.**
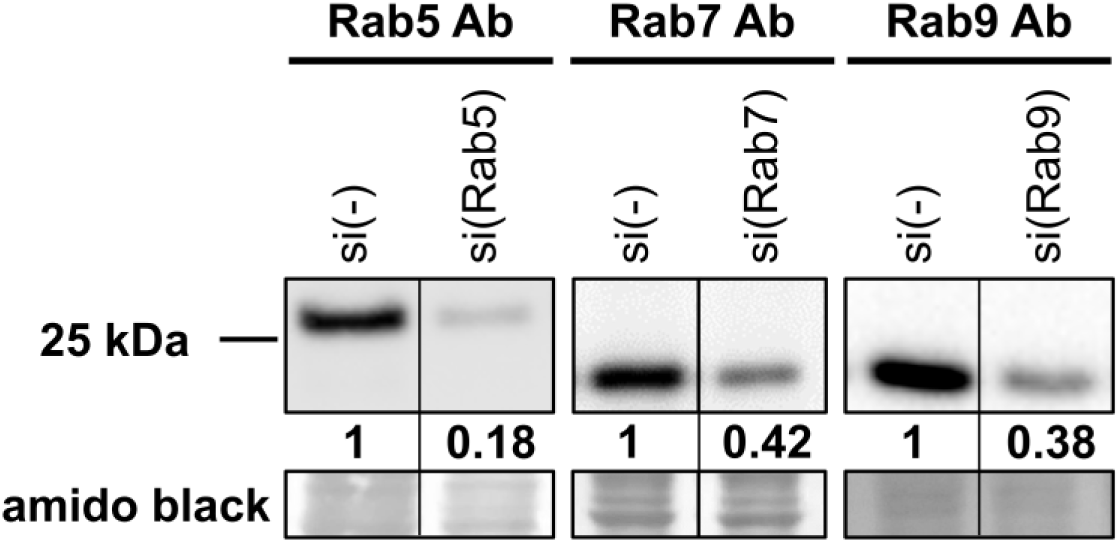
Expression of Rab5, Rab7 or Rab9 proteins in A549 cells transfected with non-specific or Rab specific siRNA. The efficiency of Rab5, Rab7 or Rab9 protein downregulation was evaluated by SDS-PAGE protein separation and western blotting using Rab specific antibodies (Ab) 48 h after transfection. The numerical values shown in the figure represent the normalized ratio of the chemiluminescent signal of the Rab5-, Rab7- or Rab9-specific bands to the total protein signal (amido black staining) relative to cells transfected with a non-specific siRNA (si(−)).

## Notes

### Competing Interest Statement

The authors have declared no competing interest.

